# Systematically gap-filling the genome-scale metabolic model of CHO cells

**DOI:** 10.1101/2020.01.27.921296

**Authors:** Hamideh Fouladiha, Sayed-Amir Marashi, Shangzhong Li, Zerong Li, Helen O. Masson, Behrouz Vaziri, Nathan E. Lewis

**Author notes:** Correspondence: Sayed-Amir Marashi, Department of Biotechnology, University of Tehran, Enghelab Avenue, Tehran 1417614411, Iran, E-mail addresses.

## Abstract

**Objective:** Chinese hamster ovary (CHO) cells are the leading cell factories for producing recombinant proteins in the biopharmaceutical industry. In this regard, constraint-based metabolic models are useful platforms to perform computational analysis of cell metabolism. These models need to be regularly updated in order to include the latest biochemical data of the cells, and to increase their predictive power. Here, we provide an update to *i*CHO1766, the metabolic model of CHO cells.

**Results:** We expanded the existing model of Chinese hamster metabolism with the help of four gap-filling approaches, leading to the addition of 773 new reactions and 335 new genes. We incorporated these into an updated genome-scale metabolic network model of CHO cells, named *i*CHO2101. In this updated model, the number of reactions and pathways capable of carrying flux is substantially increased.

**Conclusions:** The present CHO model is an important step towards more complete metabolic models of CHO cells.

## Introduction

A genome-scale metabolic network model (GEMs) is a mathematical formulation that summarizes all data about genes, proteins, and reactions known to be involved in the metabolism of a specific cell. Using reliable metabolic models, one can perform virtual (*in silico*) experiments in a rapid and inexpensive manner (Fouladiha and Marashi 2017; Gu et al. 2019). Therefore, GEMs can be helpful tools in cell biology and metabolic engineering by predicting the metabolic state of cells under certain growth conditions (Zhang and Hua 2016).

Chinese hamster ovary (CHO) cells are the main workhorse in the biopharmaceutical industry for producing recombinant proteins, such as humanized monoclonal antibodies. These cells were originally obtained from a Chinese hamster (*Cricetulus griseus*) in 1957. Several studies have focused on the optimization of the production of CHO cells using cellular and metabolic engineering methods (Wells and Robinson 2017). Experimental manipulation and maintenance of CHO cells, like many other mammalian cell lines, are costly and time-consuming. A reliable metabolic model of CHO cells can be used as a platform to perform computational analyses of cell metabolism to aid in experimental design. Such a model-driven analysis may predict the outcome of experimental tests and reduce the possibility of having false experimental results. Moreover, a CHO metabolic model can be helpful in suggesting genetic engineering and media-design strategies for improving recombinant protein production (Calmels et al. 2019; Fouladiha et al. 2020; Traustason et al. 2019). Another appreciated application of metabolic models is their role in interpreting “omics” data (Hyduke et al. 2013; Kildegaard et al. 2013; Lakshmanan et al. 2019; Richelle et al. 2019a). For example, transcriptomic and proteomic data can be mapped onto the models to infer new knowledge about the physiological characteristics of cells (Richelle et al. 2019b; Schaub et al. 2011).

One major challenge in the development of genome-scale metabolic network models is our limited knowledge of a cell’s metabolism. Specifically, genome-scale metabolic network reconstructions must be iteratively expanded as novel data emerges on enzymes and reactions that occur in the cell of interest. For example, several updates of the GEMs of *Saccharomyces cerevisia* have been published (Castillo et al. 2019), from *iND*750 (Duarte et al. 2004) and *i*IN800 (Nookaew et al. 2008), to Yeast 5 (Heavner et al. 2012), and ecYeast7 (Sánchez et al. 2017). A variety of algorithms have also been developed to predict additional reactions and potential genes that could catalyze such reactions (Karlsen et al. 2018), where using machine-learning methods have been helpful (Medlock et al. 2020; Medlock and Papin 2020). These algorithms are particularly useful for expanding the metabolic networks of non-model organisms (Biggs and Papin 2017).

The previous version of the CHO model, *i*CHO1766, has been used in several studies. For example, *i*CHO1766 was used to predict the lethality of CHO genes (Ley et al. 2019), to improve the predictive power of the model by modifying flux analysis (Chen et al. 2019; Lularevic et al. 2019), to assess heterogeneity in cell culture (Fernandez-de-Cossio-Diaz and Mulet 2019), and to improve bio-production capability of CHO cell by designing cell feeds (Fouladiha et al. 2020; Schinn et al. 2020). *i*CHO1766 has also been a helpful tool in studying metabolism of the cells, together with fluxomics (Hong et al. 2020), transcriptomics (Zhuangrong and Seongkyu 2020), and proteomics (Zhuangrong and Seongkyu 2020). In order to have more reliable and accurate results, especially in “omics” data integration, the metabolic model needs to be regularly updated to cover the latest molecular and biochemical knowledge (Schinn et al. 2020; Yeo et al. 2020).

Here, we have conducted an in-depth gap-filling of the genome-scale metabolic network reconstruction of the Chinese hamster, *i*CHO1766 (Hefzi et al. 2016), and introduce *i*CHO2101, an updated version for enhanced genome-scale modeling of CHO cell metabolism. Compared to the previous version of the CHO model, the number of genes and reactions has been increased, and the numbers of blocked reactions and dead-end metabolites have been reduced by about 10% and 15%, respectively. In other words, more parts of the metabolic model can be active, and more reactions are able to carry fluxes in this new version. These improvements increase the accuracy and precision of the predictions made by the analysis of the metabolic model.

## Methods

### Analysis of *i*CHO1766

The COBRA toolbox (Becker et al. 2007) was used for the constraint-based analysis of the metabolic model of CHO cells (*i*CHO1766). Flux Variability Analysis (FVA) (Burgard et al. 2001) was used to find the possible bounds of every flux in steady-state conditions, with no constraints on the flux bounds. If the lower and upper bounds of a specific flux were both equal to zero, that reaction was assumed to be blocked. In the same way, if the upper and lower bound of the exchange flux of a metabolite was zero, that metabolite was considered as a “non-producible and non-consumable” or a “dead-end” metabolite.

### Filling the gaps and validation of the results

In the present study, four independent approaches were used for the gap-filling of *i*CHO1766. The first two approaches were based on automatic gap-filling tools, namely, GapFind/GapFill (Kumar et al. 2007) and GAUGE (Hosseini and Marashi 2017). The GapFind algorithm uses mixed integer linear programming (MILP) to find all metabolites that cannot be produced in steady-state. The “root” gaps are those non-producible metabolites whose filling will unblock the other non-producible (or, “downstream”) gaps. Then, the GapFill algorithm selects a minimal subset of reactions from a universal reaction database that must be added to the model in order to convert a non-producible metabolite to a producible one.

In the second approach, we used GAUGE as our computational tool. GAUGE uses transcriptomics data to determine the inconsistencies between genes co-expression and flux coupling in a metabolic model. Then, GAUGE finds a minimal subset of reactions in the KEGG database whose addition can resolve the inconsistencies.

Reactions suggested by GapFind/GapFill and GAUGE (and their associated genes/proteins) were validated before being added to *i*CHO1766 as follows. If the gene ID of the new reaction or the gene ID that is attributed to the enzyme of the new reaction is found in Chinese hamster according to the KEGG database, that new reaction is confirmed. Otherwise, the validation is performed based on the results of BLASTp against the *Cricetulus griseus* (Chinese hamster) transcriptome, using the enzyme of the new reactions and CHO cell transcribed genomic sequences. For each enzyme, in the KEGG database, the amino acid sequences from different species were examined, and the best BLASTp hit was reported. A gene/protein was assumed to be present in Chinese hamster metabolism if a BLAST search hit is found with *e*-value < 1×10^−10^. To have a stricter standard, we only considered hits with query coverage >70%, or, those hits which were of >30% sequence similarity.

Our third gap-filling approach was based on manual assessment of the blocked reactions in *i*CHO1766. In several cases, the absence of an exchange or transport reaction was the cause of reaction blockage in the model. In such cases, we checked if each non-producible or non-consumable metabolite is reported in the Human Metabolome Database (HMDB) (Wishart et al. 2017). If the blocked metabolite was reported to be present in any of the human biofluids (including blood, saliva, and urine), it was assumed that the transport of the metabolite across extracellular membrane of a typical mammalian cell is possible, and therefore, an exchange reaction of that metabolite was added to the model with a high confidence score. If a metabolite was “expected” to be present in biofluids by HMDB, the exchange reaction of that metabolite was added to the model with a low confidence score.

In the fourth approach, the BiGG database (King et al. 2015) was used to retrieve all known biochemical reactions and their corresponding enzymes. Then, the KEGG database was queried to extract the full list of Chinese hamster genes and their association with biochemical enzymes. The intersection of these two lists was considered as the list of potential reactions. Then, the 1766 genes that were present in *i*CHO1766 were subtracted from the list of potential reactions to find those CHO reactions that have counterparts in BiGG, but are not present in *i*CHO1766.

### Analysis of *i*CHO2101

The COBRA toolbox (Becker et al. 2007) was used for performing flux balance analysis (FBA) and flux variability analysis (FVA) of the updated CHO model when uptake fluxes were unconstrained/constrained. In the unconstrained state, no restrictions were applied to the flux bounds. In the constrained state, on the other hand, only the metabolites of the cell culture medium were allowed to be imported to the model, with a limited flux as defined in *i*CHO1766 (Hefzi et al. 2016). Here, FBA was used to predict the maximum growth rate, and FVA was used to calculate possible flux bounds of each reaction while maintaining the maximum growth rate. The reactions with non-zero flux bounds in FVA were considered as “active” reactions.

### Gene expression analysis

In order to evaluate the new version of the model and compare it with *i*CHO1766, the transcriptomic data of CHO cells were used. These normalized data include expression levels of more than 23000 genes of CHO-S and CHO-K1 cells across 191 different samples, including published data (Hefzi et al. 2016; Van Wijk et al. 2017) and unpublished data sets. Data were processed as follows: FastQC v11.1 (Andrews 2010) was used to assess read quality. Trimmomatic v0.33 (Bolger et al. 2014) was used to trim reads with adapters or low-quality scores. STAR2.4.2a (Dobin et al. 2013) was used to align trimmed reads to the CHO-K1 genome (Xu et al. 2011), followed by calculating fpkm using cufflinks (v2.2.1).

To represent the expression of each gene, the average expression was computed across all 191 samples. The expression of a single-gene reaction was assumed to be proportional to its gene expression. In case of reactions associated with multiple genes, we restricted our analysis to those reactions whose genes were linked either with “OR” or “AND”. If all genes of a reaction were linked by “OR”/”AND”, the maximum/minimum amounts of gene expressions were attributed to that reaction. Then, we assessed expressions of the reactions in a pathway and compared it with the percentage of blocked reactions in that pathway.

## Results

### A quarter of reactions in *i*CHO1766 are blocked

The community-consensus genome-scale metabolic models of CHO cells, *i*CHO1766, includes 1766 genes, 6663 reactions, and 4455 metabolites. Using constraint-based modeling (see Methods), one can observe that about 23% of the reactions (1503 reactions out of the total 6663 reactions) of *i*CHO1766 are blocked. These blocked reactions cannot carry a non-zero flux in steady-state conditions. The reactions of *i*CHO1766 are categorized in 125 metabolic pathways, of which 83 pathways include ten or more reactions. Among these, there are 16 pathways in which at least 50% of the reactions are blocked (Table 1). The distribution of blocked reactions in all metabolic pathways has been shown in Supplementary Table 1. In addition, about 21% of the metabolites (955 metabolites out of total 4455 metabolites) in *i*CHO1766 are “dead-end” metabolites, *i.e.*, they cannot be produced nor consumed in steady-state. These metabolites belong to different subcellular parts of the model (Table 2).

**Table 1.** A list of metabolic pathways of *i*CHO1766 that more than 50 percent of the metabolic reactions in that pathway is blocked.

| Biochemical Pathway | Number of Blocked Reactions | Total Number of Reactions | Percent Blocked Reactions (%) |
| --- | --- | --- | --- |
| Chondroitin synthesis | 45 | 45 | 100 |
| Linoleate metabolism | 14 | 14 | 100 |
| Selenoamino acid metabolism | 16 | 16 | 100 |
| Vitamin E metabolism | 23 | 23 | 100 |
| Xenobiotics metabolism | 25 | 25 | 100 |
| Arachidonic acid metabolism | 72 | 73 | 98.63 |
| Androgen and estrogen synthesis and metabolism | 49 | 50 | 98 |
| Eicosanoid metabolism | 212 | 244 | 86.89 |
| N-glycan biosynthesis | 64 | 77 | 83.12 |
| Vitamin D metabolism | 22 | 29 | 75.86 |
| Miscellaneous | 45 | 69 | 65.22 |
| Vitamin c metabolism | 8 | 14 | 57.14 |
| Tyrosine metabolism | 59 | 106 | 55.66 |
| Tryptophan metabolism | 36 | 66 | 54.55 |
| Glycosphingolipid metabolism | 7 | 13 | 53.85 |
| Urea cycle | 33 | 63 | 52.38 |

**Table 2.** The distribution of dead-end metabolites of *i*CHO1766 in each subcellular part.

| Subcellular part | Total number of metabolites | Number of the blocked metabolites | Percent of the dead-end metabolites (%) |
| --- | --- | --- | --- |
| Extracellular [e] | 606 | 56 | 9.24 |
| Cytoplasm [c] | 1652 | 323 | 19.55 |
| **Endoplasmic reticulum [r]** | 479 | 153 | 31.94 |
| **Mitochondrion [m]** | 620 | 171 | 27.58 |
| **Peroxisome [x]** | 318 | 133 | 41.82 |
| **Nucleus [n]** | 158 | 66 | 41.77 |
| **Lysosome [l]** | 260 | 33 | 12.69 |
| **Golgi apparatus [g]** | 361 | 20 | 5.54 |

These blocked reactions and dead-end metabolites suggest that *i*CHO1766 includes metabolic gaps (Orth and Palsson 2010), which is common in genome-scale metabolic models. Other gaps may also exist in the model, all of which may result in the inconsistencies between model predictions and experimental results. In other words, gaps may decrease the reliability of phenotypic predictions of a metabolic model. Several gap-filling methods have been designed to find the gaps and predict the ways of removing them from the model (Pan and Reed 2018). The majority of these methods use a comprehensive dataset of all known biochemical reactions, which is often obtained from the KEGG database (Kanehisa et al. 2016). These methods try to find a subset of reactions to be added to the model to fill the gaps and improve model predictions. Gap-filling methods can be classified into three groups. The first group consists of solely-computational methods, which use different computational algorithms and linear or mixed integer linear programming (MILP) to fill the gaps of a model. GapFind/GapFill (Kumar et al. 2007), BNICE (Hatzimanikatis et al. 2005), FBA-Gap (Brooks et al. 2012), MetaFlux (Latendresse et al. 2012), FastGapFill (Thiele et al. 2014), and FastGapFilling (Thiele et al. 2014) are some examples of the first group of methods. The second group of gap-filling methods is phenotype-based methods. These methods take advantage of phenotypic data of the cells, such as viability on different carbon or nitrogen sources, to acquire new data regarding the biochemical reactions of the cell and fill the gaps of the metabolic model of the cell. Smiley (Reed et al. 2006), GrowMatch (Kumar and Maranas 2009), OMNI (Herrgård et al. 2006), and MinimalExtension (Christian et al. 2009) belong to the second group. All methods that use various kinds of “omics” data to fill the gaps of a metabolic model are in the third group, *e.g.*, Sequence-based (Krumholz and Libourel 2015) and Likelihood-based (Benedict et al. 2014) methods, Mirage (Vitkin and Shlomi 2012), and GAUGE (Hosseini and Marashi 2017).

In the present study, we decided to use GapFind/GapFill and GAUGE methods to fill the gaps of *i*CHO1766. The results of these two methods were manually validated and added to the model. Besides, two manual gap-filling approaches have been used (see Methods). In the end, representing statistics of the new model and mapping gene expression data will indicate significant improvements in CHO metabolic model.

### Gap filling approaches

Two automatic approaches, namely, GapFind/GapFill and GAUGE, and two manual approaches, were used to fill the gaps of *i*CHO1766. The GapFill method suggested the addition of 121 reactions to the model in order to enable 123 metabolites to be producible (listed in Supplementary Table 2). Some of these 121 reactions can make more than one metabolite to be producible. We validated the predictions by manually searching the KEGG database and also using BLASTp. For example, 4-coumarate (C00811) was a ‘root’ gap in *i*CHO1766 (a non-producible metabolite in steady-state). In addition, caffeate (C01197) can only be produced from 4-coumarate, and therefore, caffeate was a ‘downstream’ gap. A reaction (R00737), which is catalyzed by tyrosine ammonia-lyase, can fill both of the aforementioned gaps by transforming tyrosine to 4-coumarate and ammonia. The possibility of tyrosine ammonia-lyase expression in CHO cells was approved using the BLASTp method and therefore, R00737 was added to the model. In total, the addition of 56 reactions was validated, which enabled 87 metabolites to be producible in *i*CHO1766 (Table 3). These new reactions were associated with 30 new genes, which were added to the latest version of the model.

**Table 3.**
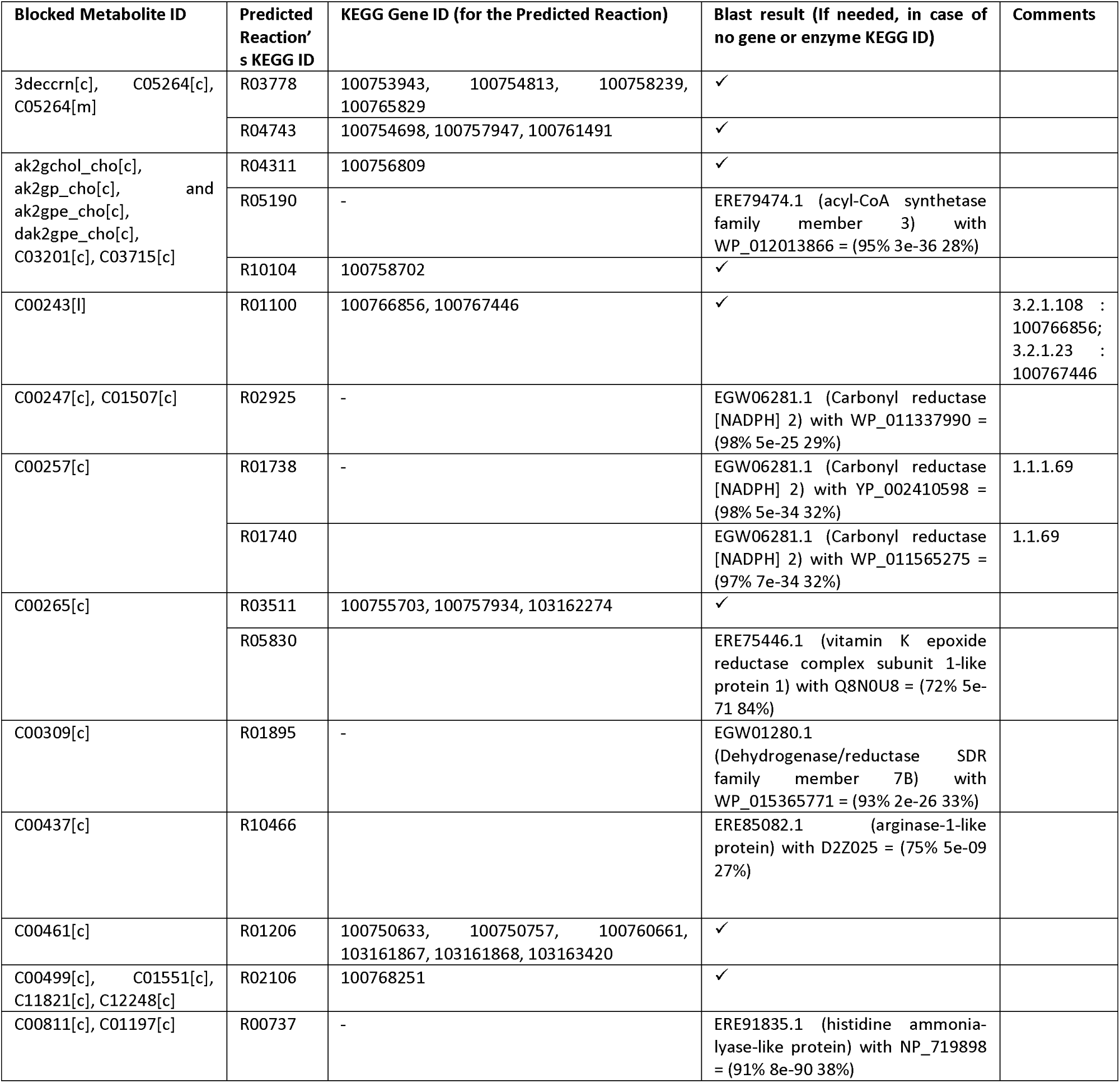
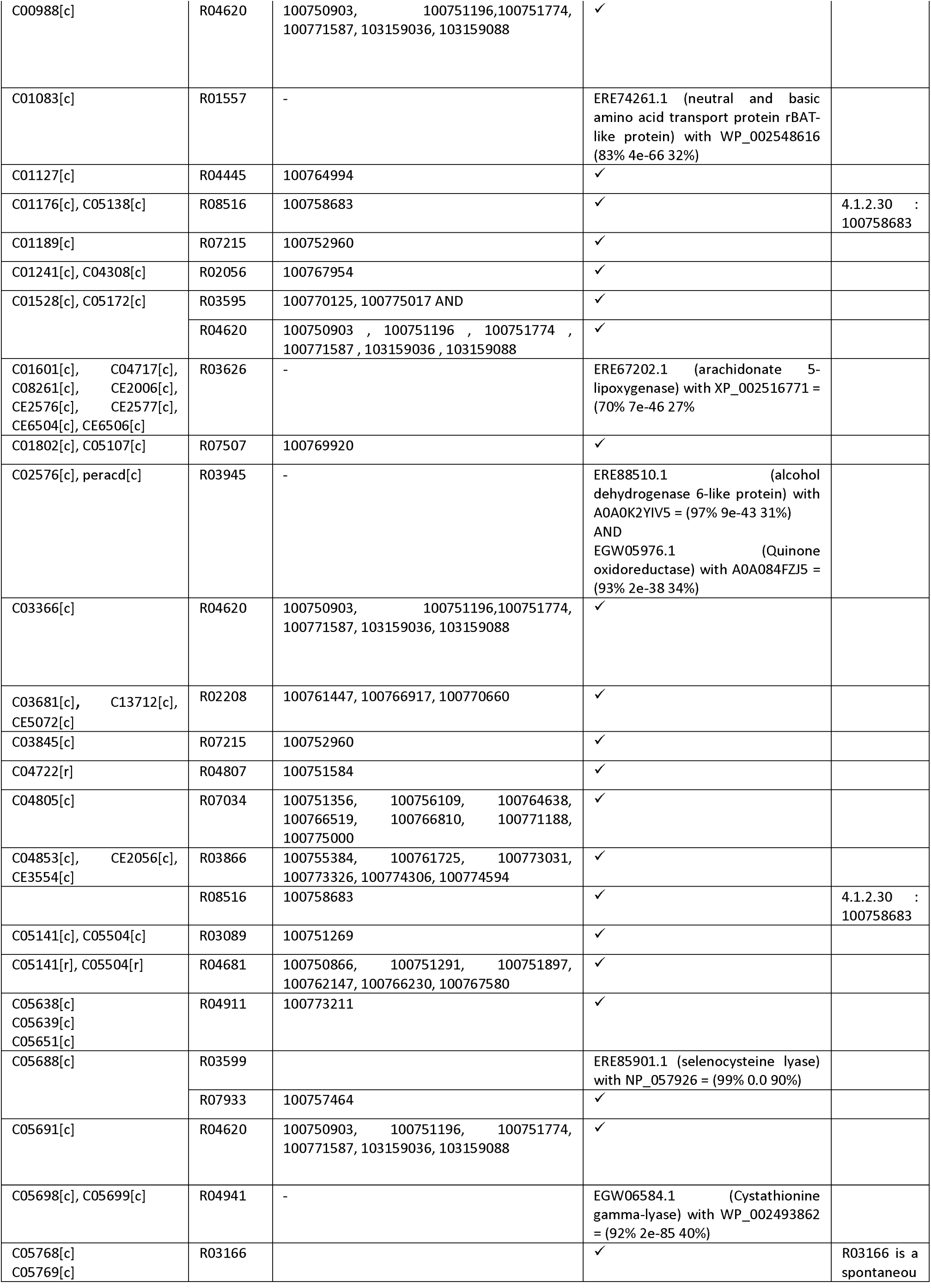
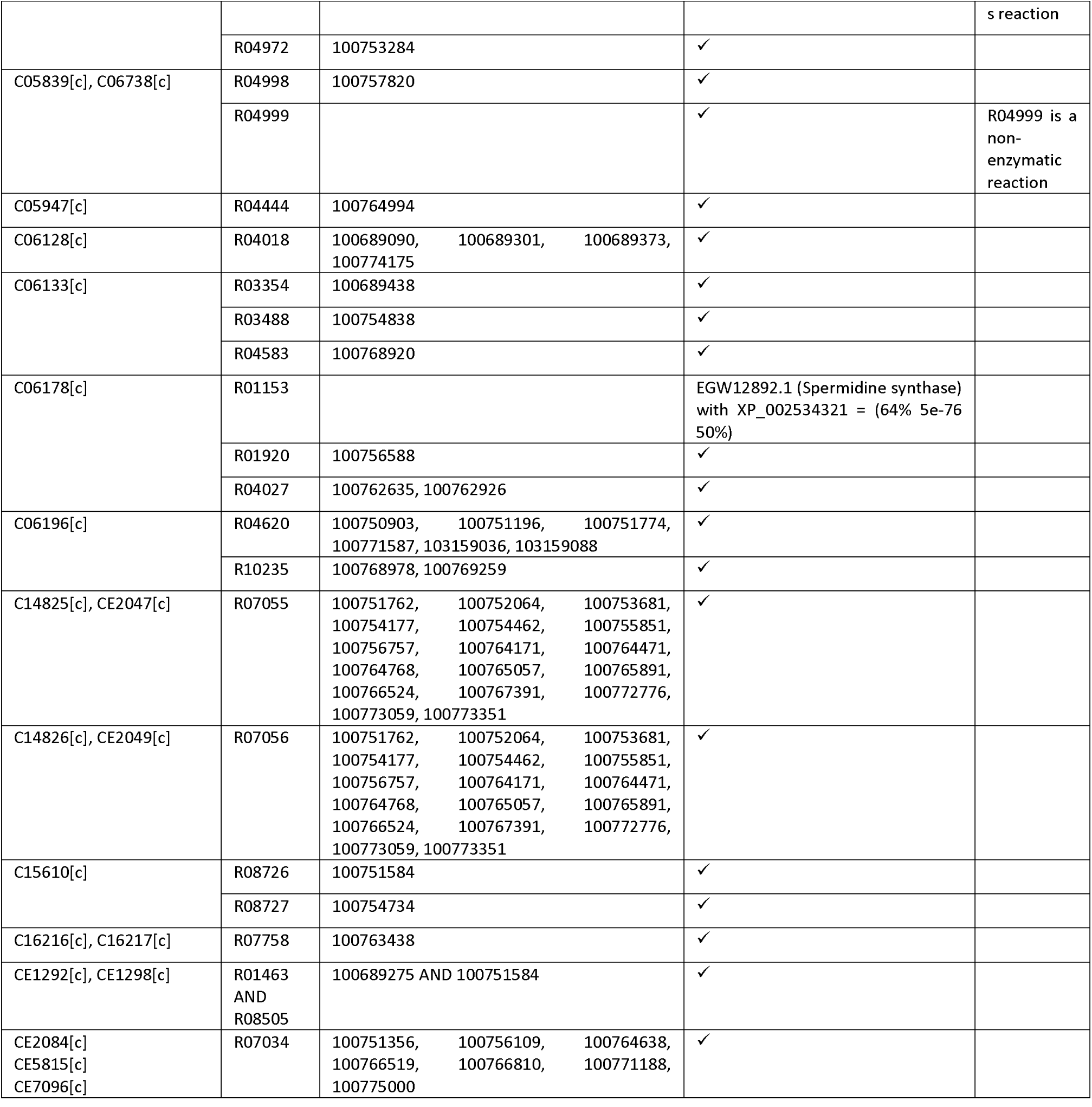
New validated reactions predicted by using the GapFill method to be added to the model. The numbers in parenthesis are query coverage, *e*-value, and sequence similarity, respectively.

Using the GAUGE method, the inconsistencies between gene co-expression and flux coupling relation of 146 gene pairs were found. GAUGE also suggested solutions for removing the inconsistencies of 64 pairs of them (listed in Supplementary Table 3). Only 37 out of 64 pairs had validated reactions as solutions. In total, 29 reactions were added to *i*CHO1766 using the GAUGE method (Table 4). These new reactions were associated with 3 new genes, which were added to the new version of the model.

**Table 4.**
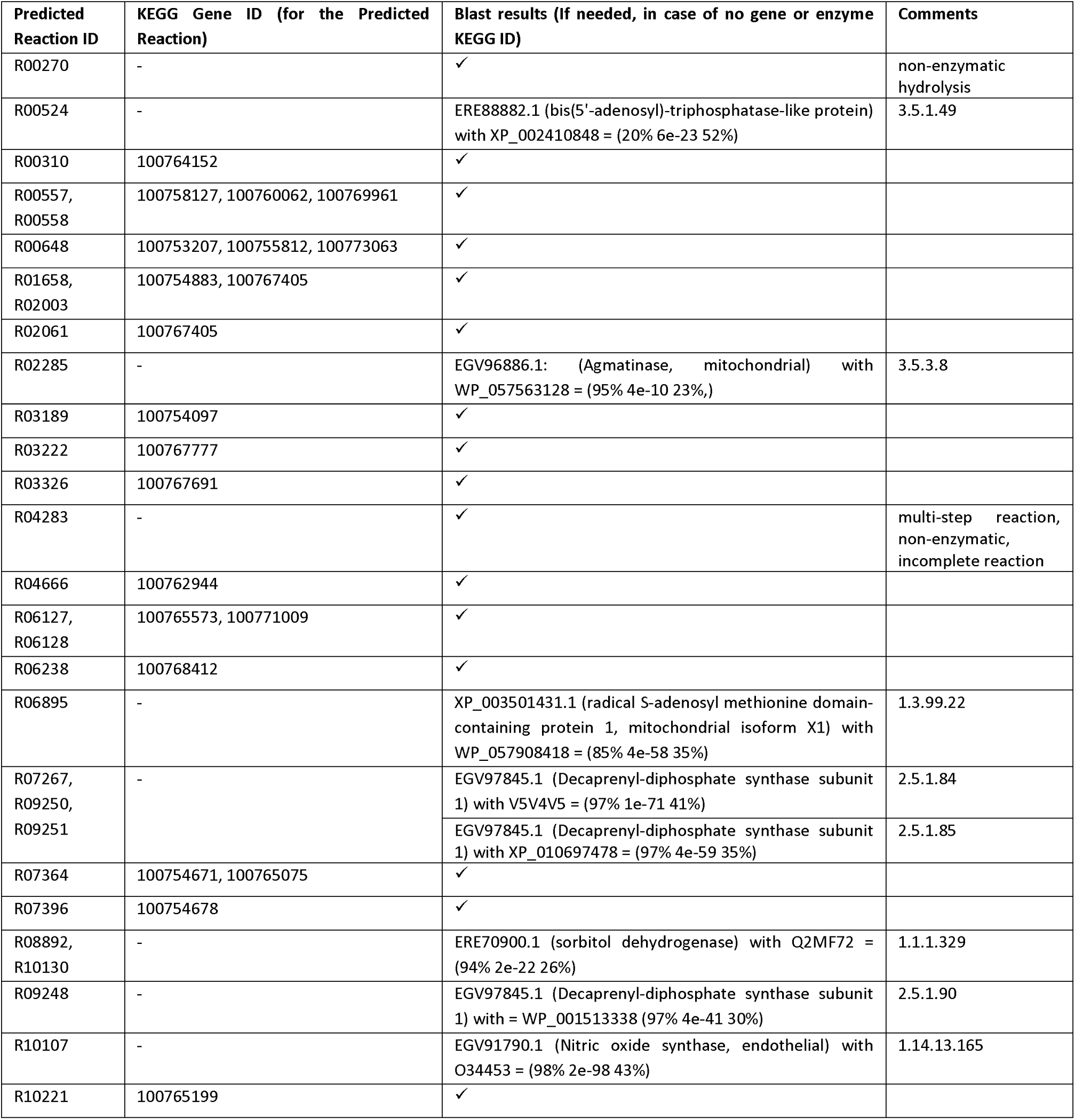
New validated reactions predicted by using the GAUGE method to be added to the model.

In the third gap-filling approach, all non-producible and non-consumable metabolites were searched in the HMDB database, and the equivalent IDs were retrieved. If any of the metabolites were detected in human biofluids, the exchange reaction of that metabolite was added to the model with a high level of confidence. This approach added 257 new reactions to the model (a full list of reactions and HMDB IDs are available in Supplementary Table 4). For example, nonanoate was a dead-end metabolite, which was detected in blood, feces, saliva, and sweat (HMDB0000847). The extracellular export of nonanoate enabled a blocked reaction to carry flux in the linoleate metabolic pathway. There was another group of metabolites that were labeled as “expected to be detected in human biofluids” by HMDB. The exchange reactions of 196 metabolites of this group were added to the model with a low level of confidence (a full list of reactions and HMDB IDs are available in Supplementary Table 5).

With a manual assessment of the blocked reactions in *i*CHO1766, we found that there was a lot of repetition of reactions in different subcellular compartments of the model. In other words, these reactions have the same reactants and products, with precisely the same stoichiometric coefficients, but in different subcellular compartments. In such cases, the absence of appropriate transport reactions caused a lot of blocked reactions. There were 178 blocked repetitive reactions in the *i*CHO1766, which have no genes, which we therefore suggest for deletion in future curation efforts (all such reactions are listed in Supplementary Table 6). Furthermore, if there was a transport reaction for a metabolite in a subcellular part with no genes in *i*CHO1766, the addition of another transport reaction for that metabolite between other subcellular parts of the new version of the model had a high confidence score. These 139 reactions were added to the new model (Supplementary Table 7).

We found 314 new genes in the fourth approach by searching the BiGG and KEGG databases (see Supplementary Table 8). Twelve of these 314 new genes were also predicted by GapFind/GapFill, and 1 out of 314 new genes was also predicted by GAUGE. The addition of these new genes updated the gene association data of 30 reactions of *i*CHO1766 and also caused 42 new reactions to be added to the new model.

### Analysis of *i*CHO2101

Using the four mentioned gap-filling approaches, a total number of 773 new reactions, 335 new genes, and 72 metabolites were added to *i*CHO1766. In addition, we reviewed the names of metabolites and reactions of the model and renamed the unknown IDs based on BiGG database. The new version of *i*CHO1766, which is named *i*CHO2101, has 2101 genes, 7436 reactions, and 4527 metabolites (see Supplementary Table 10). In *i*CHO2101, 58 pathways contain no blocked reactions, and only 5 pathways have more than 50% blocked reactions (Table 5). In addition, the distribution of dead-end metabolites of *i*CHO2101 in different subcellular compartments has been reduced to less than 10% (Table 6). Figure 1 summarizes the improvements made in the current study for the metabolic model of CHO cells by creating a visual comparison of model statistics, blocked reactions, and dead-end metabolites between *i*CHO1766 and *i*CHO2101.

**Table 5.** A list of metabolic pathways of *i*CHO2101 that more than 50 percent of the metabolic reactions in that pathway is blocked.

| Biochemical Pathway | Number of Blocked Reactions | Total Number of Reactions | Percent Blocked Reactions (%) |
| --- | --- | --- | --- |
| Xenobiotics metabolism | 25 | 25 | 100 |
| Selenoamino acid metabolism | 15 | 21 | 71.43 |
| Androgen and estrogen synthesis and metabolism | 29 | 51 | 56.86 |
| Arachidonic acid metabolism | 42 | 74 | 56.76 |
| Eicosanoid metabolism | 127 | 245 | 51.84 |

**Table 6.** The distribution of dead-end metabolites of *i*CHO2101 in each subcellular part.

| **Subcellular part** | **Total number of metabolites** | **Number of the dead-end metabolites** | **Percent of the dead-end metabolites (%)** |
| --- | --- | --- | --- |
| **Extracellular [e]** | 609 | 2 | 0.33 |
| **Cytoplasm [c]** | 1713 | 156 | 9.10 |
| **Endoplasmic reticulum [r]** | 482 | 45 | 9.34 |
| **Mitochondrion [m]** | 625 | 52 | 8.32 |
| **Peroxisome [x]** | 318 | 26 | 8.18 |
| **Nucleus [n]** | 159 | 12 | 7.55 |
| **Lysosome [l]** | 260 | 3 | 1.15 |
| **Golgi apparatus [g]** | 361 | 2 | 0.55 |

**Figure 1.**
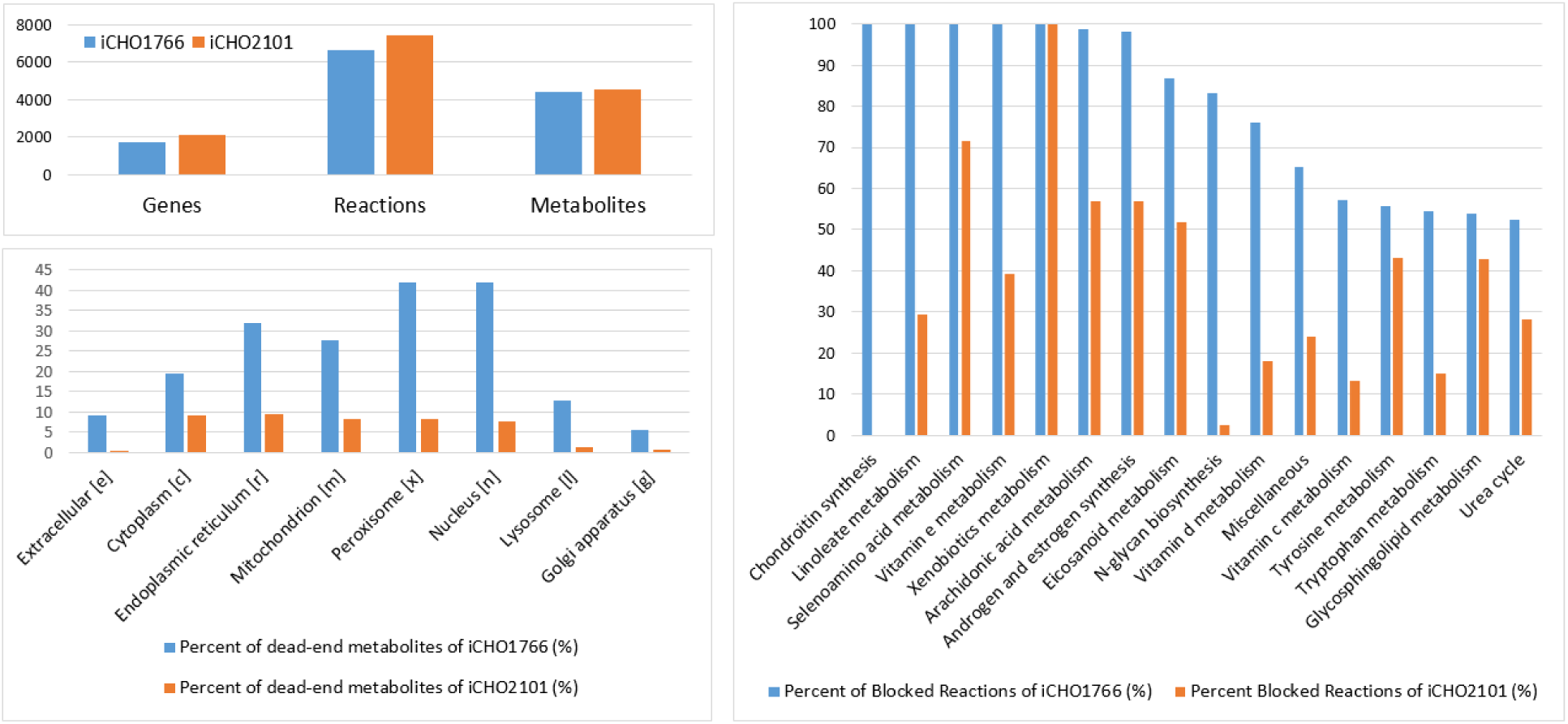
A visual comparison between the statistics of *i*CHO1766 and *i*CHO2101. Part (A) shows the number of genes, reactions, and metabolites. Part (B) shows the distribution of dead-end metabolites in different subcellular parts. Part (C) shows the percent of blockage in the selected pathways reported in Table 1.

Using FBA after applying our published uptake and secretion constraints, we found the maximum growth rate in the constrained state was similar for *i*CHO1766 and *i*CHO2101 (0.03 h^-1^). By performing FVA in the constrained state of *i*CHO1766 and *i*CHO2101, we found the number of “active” reactions in each metabolic pathway had been significantly improved in the gap-filled version of the model. Figure 2 shows the percent of activities of fluxes in 14 metabolic pathways with more than 5 reactions, where the changes between *i*CHO1766 and *i*CHO2101 are more than 30%. For example, all reactions of ‘sphingolipid metabolism’ are “active” in modeling the growth using *i*CHO2101, thus enabling the analysis of this process, which has been previously reported to be of importance for the growth of CHO cells (Hanada et al. 1992).

**Figure 2.**
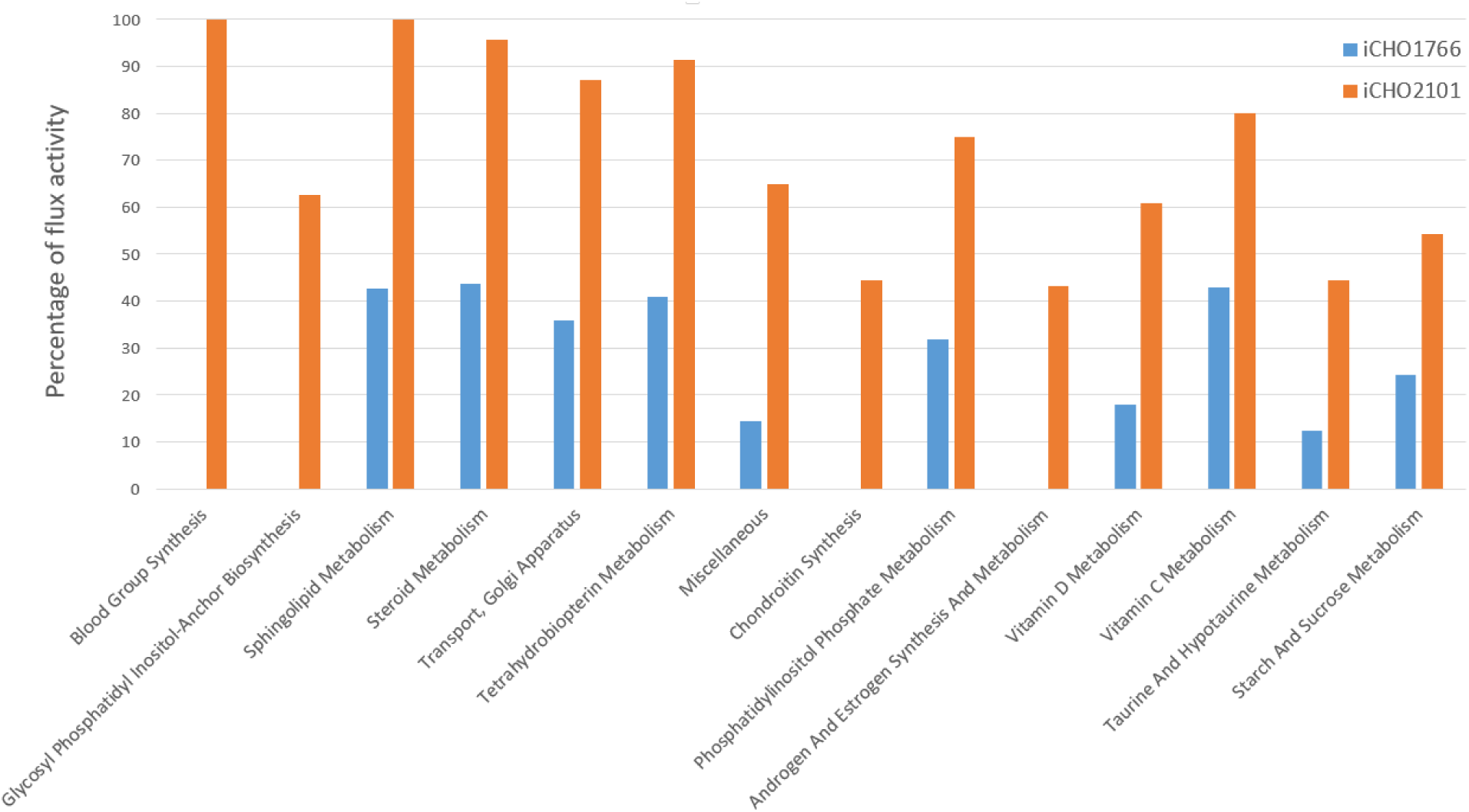
A visual comparison of flux activities in 14 metabolic pathways of *i*CHO1766 and *i*CHO2101, where the changes are more than 30% in comparison.

### Gene expression analysis

We subsequently analyzed the expression of the genes in the metabolic models in 191 RNA-Seq samples. We computed the expression levels of reactions (see Methods). Then, considering the expressions of reactions in the metabolic pathways of the *i*CHO1766, it was revealed that some of the pathways with a high level of expression had a high percent of blockage. For example, ‘androgen and estrogen synthesis and metabolism’ had the highest level of expression among blocked pathways, where 98% of the reactions were blocked. In the new model, only 56% of the reactions in the mentioned pathway are still blocked. In another example, ‘glyoxylate and dicarboxylate metabolism,’ ‘methionine and cysteine metabolism,’ and ‘galactose metabolism’ are among the top ten highly expressed pathways, while about 30% of the reactions are blocked in the pathways in *i*CHO1766. In *i*CHO2101, the blocked reactions of the three pathways have been reduced to 11%, 15%, and 7%, respectively. A full list of the pathways and expression levels is available in Supplementary Table 9.

## Discussion

In the present study, four approaches were used to fill the gaps of *i*CHO1766. At first, we used GapFill that successfully filled 12% (124 out of 1049) of no-production metabolites. Then, using GAUGE, 40% (28 out of 71) of the inconsistencies between genes co-expression and flux coupling relations of reaction pairs were fixed. Furthermore, exchange and transport reactions of the model were revised, using HMDB database. Finally, new genes were added to the model based on KEGG and BiGG databases. All newly predicted reactions and metabolites were subsequently added to the model to generate a new version of the CHO metabolic model, named *i*CHO2101. In total, the percentage of blocked reactions was 21.6% (1441 out of 6663) in *i*CHO1766, which has been reduced to 11.3% (837 out of 7336) in *i*CHO2101. In addition, the percentage of dead-end metabolites from 21.4% (955 out of 4456) in *i*CHO1766 has been reduced to 6.6% (298 out of 4527) in *i*CHO2101. The addition of these new reactions, metabolites, and genes can increase the scope of pathways that can be simulated in CHO cells, and increase the reliability of the model predictions in general for CHO cells with more comprehensive models of CHO cell metabolism.

The importance of CHO cells in the pharmaceutical industry producing recombinant protein drugs is evident. In this regard, due to the notable drawbacks of the present kinetic models (Carinhas et al. 2012), a constraint-based metabolic model can be beneficial to have an *in silico* platform to mechanistically model the metabolism of CHO cells. For example, the limiting factors of cell culture can be easily modelled by constraining the exchange fluxes of the model. In addition, integration of “omics” data with a constraint-based metabolic model can shed light on the metabolism of CHO cells.

Bioprocess optimization of CHO cells has been a major topic of research, including studies which focused on the design of compositions of cell culture media (Galbraith et al. 2018; Ritacco et al. 2018). Mammalian cell culture media are mostly composed of amino acids. Amino acid metabolism greatly influences the viability and production of CHO cells (Salazar et al. 2016). The average percentage of blocked reactions in the metabolic pathways of different amino acids was reduced from 34.10% in *i*CHO1766 to 13.56% in *i*CHO2101. Therefore, the applicability of CHO model in bioprocess studies can be increased by refining the metabolic models. Recently, an extended version of the GEM of CHO cells was released, in which new constraints were added to the model based on enzyme capacity of the reactions (Yeo et al. 2020). Yet, the focus of our study is to fill the gaps and manually curate the previous model (Hefzi et al. 2016). In conclusion, with more active metabolic pathways and more precise gene-protein-reaction associations in a GEM of CHO cells, one is able to infer more accurate cell line-specific models. Such models can address the cell-specific metabolic signatures of different cell lines for better predicting biopharmaceutical production capabilities (Carinhas et al. 2013).

## Acknowledgments

This work was facilitated through generous funding from the Novo Nordisk Foundation through Center for Biosustainability at the Technical University of Denmark (NNF10CC1016517).

## Supplementary Information

Supplementary Table 1: The distribution of blocked reactions of *i*CHO1766 in all metabolic pathways.

Supplementary Table 2: The list of new reactions added by using GapFill method.

Supplementary Table 3: The list of new reactions added by using GAUGE method.

Supplementary Table 4: The list of metabolites that were labeled as “detected in human biofluids” in HMDB and the new reactions associated with them.

Supplementary Table 5: The list of metabolites that were labeled as “expected to be detected in human biofluids” in HMDB and the new reactions associated with them.

Supplementary Table 6: The list of blocked repetitive reactions in *i*CHO1766 that have been suggested for deletion.

Supplementary Table 7: The list of new transport reactions that have a similar reaction in a subcellular part with no genes in *i*CHO1766.

Supplementary Table 8: The list of new reactions added by searching the BiGG database.

Supplementary Table 9: The list of the reactions in metabolic pathways and expression levels associated with them, both in *i*CHO1766 and *i*CHO2101.

Supplementary Table 10: The spreadsheet format of *i*CHO2101.

## Compliance with ethical standards

### Ethical approval

This study does not include any studies with human participants or animals performed by any of the authors

## References

Andrews S (2010) FastQC: a quality control tool for high throughput sequence data. Babraham Bioinformatics, Babraham Institute, Cambridge, United Kingdom.

Becker SA, Feist AM, Mo ML, Hannum G, Palsson BØ, Herrgard MJ (2007) Quantitative prediction of cellular metabolism with constraint-based models: the COBRA Toolbox. Nat Protoc 2:727–738

Benedict MN, Mundy MB, Henry CS, Chia N, Price ND (2014) Likelihood-based gene annotations for gap filling and quality assessment in genome-scale metabolic models. PLoS Comput Biol 10:e1003882

Biggs MB, Papin JA (2017) Managing uncertainty in metabolic network structure and improving predictions using EnsembleFBA. PLoS Comput Biol 13:e1005413

Bolger AM, Lohse M, Usadel B (2014) Trimmomatic: a flexible trimmer for Illumina sequence data. Bioinformatics 30:2114–2120

Brooks JP, Burns WP, Fong SS, Gowen CM, Roberts SB (2012) Gap detection for genome-scale constraint-based models. Advances in bioinformatics 2012: 323472

Burgard AP, Vaidyaraman S, Maranas CD (2001) Minimal reaction sets for *Escherichia coli* metabolism under different growth requirements and uptake environments. Biotechnol Prog 17:791–797

Calmels C, McCann A, Malphettes L, Andersen MR (2019) Application of a curated genome-scale metabolic model of CHO DG44 to an industrial fed-batch process. Metab Eng 51:9–19

Carinhas N, Duarte TM, Barreiro LC, Carrondo MJ, Alves PM, Teixeira AP (2013) Metabolic signatures of GSDCHO cell clones associated with butyrate treatment and culture phase transition. Biotechnol Bioeng 110:3244–3257

Carinhas N, Oliveira R, Alves PM, Carrondo MJ, Teixeira AP (2012) Systems biotechnology of animal cells: the road to prediction. Trends Biotechnol 30:377–385

Castillo S, Patil KR, Jouhten P (2019) Yeast genome-scale metabolic models for simulating genotype– phenotype relations. In: Yeasts in Biotechnology and Human Health. Springer, pp 111–133

Chen Y, McConnell BO, Dhara VG, Naik HM, Li C-T, Antoniewicz MR, Betenbaugh MJ (2019) An unconventional uptake rate objective function approach enhances applicability of genome-scale models for mammalian cells. NPJ Syst Biol Appl 5:1–11

Christian N, May P, Kempa S, Handorf T, Ebenhöh O (2009) An integrative approach towards completing genome-scale metabolic networks. Mol BioSyst 5:1889–1903

Dobin A et al. (2013) STAR: ultrafast universal RNA-seq aligner. Bioinformatics 29:15–21

Duarte NC, Herrgård MJ, Palsson BØ (2004) Reconstruction and validation of *Saccharomyces cerevisiae* iND750, a fully compartmentalized genome-scale metabolic model. Genome Res 14:1298–1309

Fernandez-de-Cossio-Diaz J, Mulet R (2019) Maximum entropy and population heterogeneity in continuous cell cultures. PLoS Comput Biol 15:e1006823

Fouladiha H, Marashi S-A (2017) Biomedical applications of cell-and tissue-specific metabolic network models. J Biomed Inform 68:35–49

Fouladiha H, Marashi S-A, Torkashvand F, Mahboudi F, Lewis NE, Vaziri B (2020) A metabolic network-based approach for developing feeding strategies for CHO cells to increase monoclonal antibody production. Bioprocess Biosyst Eng 43:1381–1389

Galbraith SC, Bhatia H, Liu H, Yoon S (2018) Media formulation optimization: current and future opportunities. Curr Opin Chem Eng 22:42–47

Gu C, Kim GB, Kim WJ, Kim HU, Lee SY (2019) Current status and applications of genome-scale metabolic models. Genome Biol 20:121

Hanada K, Nishijima M, Kiso M, Hasegawa A, Fujita S, Ogawa T, Akamatsu Y (1992) Sphingolipids are essential for the growth of Chinese hamster ovary cells. Restoration of the growth of a mutant defective in sphingoid base biosynthesis by exogenous sphingolipids. J Biol Chem 267:23527–23533

Hatzimanikatis V, Li C, Ionita JA, Henry CS, Jankowski MD, Broadbelt LJ (2005) Exploring the diversity of complex metabolic networks. Bioinformatics 21:1603–1609

Heavner BD, Smallbone K, Barker B, Mendes P, Walker LP (2012) Yeast 5–an expanded reconstruction of the *Saccharomyces cerevisiae* metabolic network. BMC Syst Biol 6:55

Hefzi H et al. (2016) A consensus genome-scale reconstruction of Chinese hamster ovary cell metabolism. Cell Syst 3:434–443

Herrgård MJ, Fong SS, Palsson BØ (2006) Identification of genome-scale metabolic network models using experimentally measured flux profiles. PLoS Comput Biol 2:e72

Hong JK, Yeo HC, Lakshmanan M, Han S-h, Cha HM, Han M, Lee D-Y (2020) *In silico* model-based characterization of metabolic response to harsh sparging stress in fed-batch CHO cell cultures. J Biotechnol 308:10–20

Hosseini Z, Marashi S-A (2017) Discovering missing reactions of metabolic networks by using gene coexpression data. Sci Rep 7:41774

Hyduke DR, Lewis NE, Palsson BØ (2013) Analysis of omics data with genome-scale models of metabolism. Mol BioSyst 9:167–174

Kanehisa M, Furumichi M, Tanabe M, Sato Y, Morishima K (2016) KEGG: new perspectives on genomes, pathways, diseases and drugs. Nucleic Acids Res 45:D353–D361

Karlsen E, Schulz C, Almaas E (2018) Automated generation of genome-scale metabolic draft reconstructions based on KEGG. BMC bioinformatics 19:467

Kildegaard HF, Baycin-Hizal D, Lewis NE, Betenbaugh MJ (2013) The emerging CHO systems biology era: harnessing the omics revolution for biotechnology. Curr Opin Biotechnol 24:1102–1107

King ZA et al. (2015) BiGG Models: A platform for integrating, standardizing and sharing genome-scale models. Nucleic Acids Res 44:D515–D522

Krumholz EW, Libourel IG (2015) Sequence-based network completion reveals the integrality of missing reactions in metabolic networks. J Biol Chem 290:19197–19207

Kumar VS, Dasika MS, Maranas CD (2007) Optimization based automated curation of metabolic reconstructions. BMC bioinformatics 8:212

Kumar VS, Maranas CD (2009) GrowMatch: an automated method for reconciling *in silico*/*in vivo* growth predictions. PLoS Comput Biol 5:e1000308

Lakshmanan M et al. (2019) MultiDomics profiling of CHO parental hosts reveals cell lineDspecific variations in bioprocessing traits. Biotechnol Bioeng 116:2117–2129

Latendresse M, Krummenacker M, Trupp M, Karp PD (2012) Construction and completion of flux balance models from pathway databases. Bioinformatics 28:388–396

Ley D et al. (2019) Reprogramming AA catabolism in CHO cells with CRISPR/Cas9 genome editing improves cell growth and reduces byproduct secretion. Metab Eng 56:120–129

Lularevic M, Racher AJ, Jaques C, Kiparissides A (2019) Improving the accuracy of flux balance analysis through the implementation of carbon availability constraints for intracellular reactions. Biotechnol Bioeng 116:2339–2352

Medlock GL, Moutinho TJ, Papin JA (2020) Medusa: software to build and analyze ensembles of genome-scale metabolic network reconstructions. PLoS Comput Biol 16:e1007847

Medlock GL, Papin JA (2020) Guiding the refinement of biochemical knowledgebases with ensembles of metabolic networks and machine learning. Cell Syst 10:109–119

Nookaew I et al. (2008) The genome-scale metabolic model *iIN800* of *Saccharomyces cerevisiae* and its validation: a scaffold to query lipid metabolism. BMC Syst Biol 2:71

Orth JD, Palsson BØ (2010) Systematizing the generation of missing metabolic knowledge. Biotechnol Bioeng 107:403–412

Pan S, Reed JL (2018) Advances in gap-filling genome-scale metabolic models and model-driven experiments lead to novel metabolic discoveries. Curr Opin Biotechnol 51:103–108

Reed JL et al. (2006) Systems approach to refining genome annotation. PNAS 103:17480–17484

Richelle A, Chiang AW, Kuo C-C, Lewis NE (2019a) Increasing consensus of context-specific metabolic models by integrating data-inferred cell functions. PLoS Comput Biol 15:e1006867

Richelle A, Joshi C, Lewis NE (2019b) Assessing key decisions for transcriptomic data integration in biochemical networks. PLoS Comput Biol 15:e1007185

Ritacco FV, Wu Y, Khetan A (2018) Cell culture media for recombinant protein expression in Chinese hamster ovary (CHO) cells: History, key components, and optimization strategies. Biotechnol Prog 34:1407–1426

Salazar A, Keusgen M, von Hagen J (2016) Amino acids in the cultivation of mammalian cells. Amino acids 48:1161–1171

Sánchez BJ, Zhang C, Nilsson A, Lahtvee PJ, Kerkhoven EJ, Nielsen J (2017) Improving the phenotype predictions of a yeast genomeDscale metabolic model by incorporating enzymatic constraints. Mol Syst Biol 13:935

Schaub J, Clemens C, Kaufmann H, Schulz TW (2011) Advancing biopharmaceutical process development by system-level data analysis and integration of omics data. In: Genomics and Systems Biology of Mammalian Cell Culture. Springer, pp 133–163

Schinn S-M, Morrison C, Wei W, Zhang L, Lewis NE (2020) A genome-scale metabolic network model synergizes with statistical learning to predict amino acid concentrations in Chinese hamster ovary cell cultures. bioRxiv

Thiele I, Vlassis N, Fleming RM (2014) fastGapFill: efficient gap filling in metabolic networks. Bioinformatics 30:2529–2531

Traustason B, Cheeks M, Dikicioglu D (2019) Computer-aided strategies for determining the amino acid composition of medium for Chinese hamster ovary cell-based biomanufacturing platforms. Int J Mol Sci 20:5464

Van Wijk XM et al. (2017) Whole-genome sequencing of invasion-resistant cells identifies laminin α2 as a host factor for bacterial invasion. MBio 8:e02128–02116

Vitkin E, Shlomi T (2012) MIRAGE: a functional genomics-based approach for metabolic network model reconstruction and its application to cyanobacteria networks. Genome Biol 13:R111

Wells E, Robinson AS (2017) Cellular engineering for therapeutic protein production: product quality, host modification, and process improvement. Biotechnol J 12:1600105

Wishart DS et al. (2017) HMDB 4.0: the human metabolome database for 2018. Nucleic Acids Res 46:D608–D617

Xu X et al. (2011) The genomic sequence of the Chinese hamster ovary (CHO)-K1 cell line. Nat Biotechnol 29:735

Yeo HC, Hong J, Lakshmanan M, Lee D-Y (2020) Enzyme capacity-based genome scale modelling of CHO cells. Metab Eng 60:138–147

Zhang C, Hua Q (2016) Applications of genome-scale metabolic models in biotechnology and systems medicine. Front Physiol 6:413

Zhuangrong H, Seongkyu Y (2020) Identifying metabolic features and engineering targets for productivity improvement in CHO cells by integrated transcriptomics and genome-scale metabolic model. Biochem Eng J:107624

